# Sterically-Confined Rearrangements of SARS-CoV-2 Spike Protein Control Cell Invasion

**DOI:** 10.1101/2021.01.18.427189

**Authors:** Esteban Dodero-Rojas, José N. Onuchic, Paul C. Whitford

## Abstract

Severe acute respiratory syndrome coronavirus 2 (SARS-CoV-2) is highly contagious, and transmission involves a series of processes that may be targeted by vaccines and therapeutics. During transmission, host cell invasion is controlled by a large-scale conformational change of the Spike protein. This conformational rearrangement leads to membrane fusion, which creates transmembrane pores through which the viral genome is passed to the host. During Spike-protein-mediated fusion, the fusion peptides must be released from the core of the protein and associate with the host membrane. Interestingly, the Spike protein possesses many post-translational modifications, in the form of branched glycans that flank the surface of the assembly. Despite the large number of glycosylation sites, until now, the specific role of glycans during cell invasion has been unclear. Here, we propose that glycosylation is needed to provide sufficient time for the fusion peptides to reach the host membrane, otherwise the viral particle would fail to enter the host. To understand this process, an all-atom model with simplified energetics was used to perform thousands of simulations in which the protein transitions between the prefusion and postfusion conformations. These simulations indicate that the steric composition of the glycans induces a pause during the Spike protein conformational change. We additionally show that this glycan-induced delay provides a critical opportunity for the fusion peptides to capture the host cell. This previously-unrecognized role of glycans reveals how the glycosylation state can regulate infectivity of this pervasive pathogen.

## Introduction

The current COVID-19 pandemic is being driven by severe acute respiratory syndrome coronavirus 2 (SARS-CoV-2). While vaccine and treatment development will help mitigate the immediate impact of this disease, long-term strategies for its eradication will rely on an understanding of the factors that control transmission. The need to isolate the molecular constituents that govern SARS-CoV-2 dynamics is widely recognized, where the global scientific community has undergone its most rapid transformation in recent history. This unprecedented redirection of scientific inquiry has provided atomic-resolution structures of SARS-CoV-2 proteins at various stages of infection (*1–6*), as well as computational analysis of specific conformational states (*7–13*). Despite these advances, our understanding of the mechanism by which SARS-CoV-2 enters the host cell is limited.

Central to the function of SARS-CoV-2 is host-cell recognition by the Spike protein, which results in virus-host membrane fusion and transfer of the viral genome. In the active virion, the Spike protein assembly (S) is a 3-fold symmetric homo-trimer (*1*), where each protein contains *~* 1200 residues (Fig. 1). The complex is anchored to the viral capsid by a transmembrane (TM) helical bundle, while the remaining regions reside on the capsid exterior. Cleavage at the S1/S2 and S2’ sites leads to activation of the Spike protein, where the resulting subunits (S1 and S2) maintain contact through non-bonded interactions (Fig. 1A) (*14*). The receptor binding domain (RBD) in S1 can then associate with the ACE2 receptor (*4, 15*), which triggers S1 release from S2 (Fig. 1B). This allows for a dramatic structural rearrangement within S2, during which the fusion peptides must bind and recruit a host cell (Fig. 1C) (*16*).

**Figure 1:**
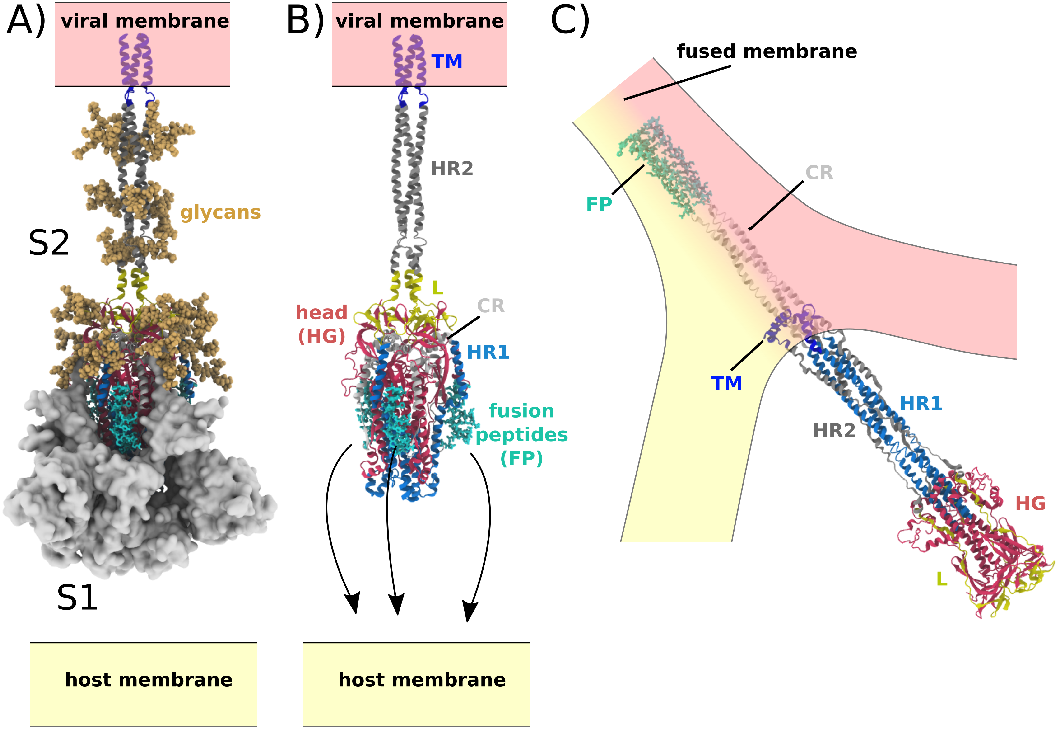
Spike-protein-mediated membrane fusion. A) The active Spike protein assembly is composed of the subunits S1 (white surface) and S2 (cartoon representation) (*2*), which remain bound through nonbonded interactions. Numerous glycosylation sites (glycans shown in orange) are present in the Head Group (HG) and Heptad Repeat 2 (HR2) regions. The modeled glycans are consistent with previous studies (*7, 13*). B) Upon recognition of the ACE2 receptor and cleavage at the S2’ site, S1 dissociates. In addition to the HG and HR2 regions, S2 is composed of the Heptad Repeat 1 (HR1), Linker (L), Fusion Peptide (FP), Connector (CR) and Transmembrane (TM) regions. Since the HR2 and TM regions were not resolved in the prefusion structure (*2*), they were modeled as a helical bundle, consistent with previous studies (*7*). C) Release of S1 allows for the FPs to associate with and recruit the host membrane. The HG and HR1 regions undergo a large-scale rotation (*>* 90^°^), leading to fusion of the host and viral membranes. Since the CR and FP regions were not resolved in the postfusion structure (*30*), they were modeled as an extended helical bundle.

Most of the current SARS-CoV-2 therapies and vaccines have focused on the ACE2 recognition stage of virus invasion. An alternate strategy is to target the conformational change in S2 that induces membrane fusion. Disruption of this process could stop, or at least impair, viral transmission. Therefore, understanding the Spike protein conformational steps during membrane fusion can provide new targets for medical applications.

The fusion process involves a global reorganization of S2 (Figs. 1 and S1). This includes dissociation of the fusion peptides (FP) from the head group (HG), disordering of Heptad Repeat 2 (HR2), rotation of the head group relative to the viral membrane and then reordering of Heptad Repeat 1 (HR1), HR2 and Connecting Region (CR) into an extended helical arrangement. During this elaborated process, the fusion peptides associate with the host membrane, where subsequent “zippering” of a HR1-HR2 superhelical structure likely provides energy to recruit the host membrane. In the postfusion structure, the TM, CR and FP regions adopt proximal positions, allowing them to facilitate membrane fusion.

While high-resolution structures have been resolved for the Spike protein in the prefusion and postfusion conformations, the precise sequence of structural changes during fusion is unknown. Accordingly, the molecular factors that control this process have yet to be determined. As an example, while many post-translational modifications (glycans) have been identified (13 and 9 on each S1 and S2 monomer) (*1, 2, 6, 17*), there has not been an investigation into their role during the fusion step of infection. However, simulations of the prefusion protein have suggested that glycans may shield the Spike protein and prevent recognition by the immune system (*7*). In addition, studies have provided evidence that glycans may serve as activators for the lectin pathway (*18, 19*). While glycans have been implicated in many aspects of the viral “life” cycle, it is not known whether they directly impact the host-entry process.

Here, we perform molecular dynamics simulations with an all-atom structure-based model to determine whether the steric composition of glycans can have a meaningful influence on SARS-CoV-2 Spike-protein-mediated membrane fusion. Simulations are initiated with the Spike protein in the prefusion conformation, while the energy landscape favors the postfusion conformation. By comparing the dynamics with and without glycans present, we show how the steric composition of the glycans can extend the lifetime of a critical intermediate in which the head appears to be sterically-caged. This leads to a transient pause that can increase the probability of successfully recruiting the host cell. These calculations provide direct physical evidence that the glycosylation state is a critical factor that determines infectivity of SARS-CoV-2.

## Results

### Simulating the membrane-fusion-associated conformational change of SARS-CoV-2 Spike protein

In order to characterize the mechanism of Spike-protein-mediated membrane fusion, we employed an all-atom structure-based model (*20, 21*) and simulated transitions between the prefusion and postfusion conformations (Fig. 2B and Movie S1). In a structure-based (Gō-like) model, some/all of the energetic interactions are defined based on knowledge of specific stable (experimentally-resolved) structures. In the context of protein folding, applying these types of models (*22*) is supported by the principle of minimal frustration (*23, 24*). However, to warrant their application to study conformational transitions, it is necessary to recognize that the models describe the effective energetics of each system (*25–27*). That is, by explicitly defining the molecular interactions to stabilize the endpoint conformations, the models are intended to provide a first-order approximation to the energetics. In the presented model, only interactions that are specific to the prefusion and postfusion configurations were defined to be stable. For the TM region, stabilizing prefusion-specific interactions allow it to serve as an anchor between the Spike protein and the viral membrane. An implicit membrane potential was also introduced to restrain the TM to a plane. Finally, all non-TM interactions were defined to stabilize the postfusion conformation (Fig. 1). Qualitatively, this model describes the Spike protein as a loaded (non-linear) spring that is released upon cleavage of the S2’ site and dissociation of S1 (Fig. 2A). While the potential energy in the model is downhill, molecular sterics can still lead to pronounced free-energy barriers that control the kinetics (*28, 29*).

**Figure 2:**
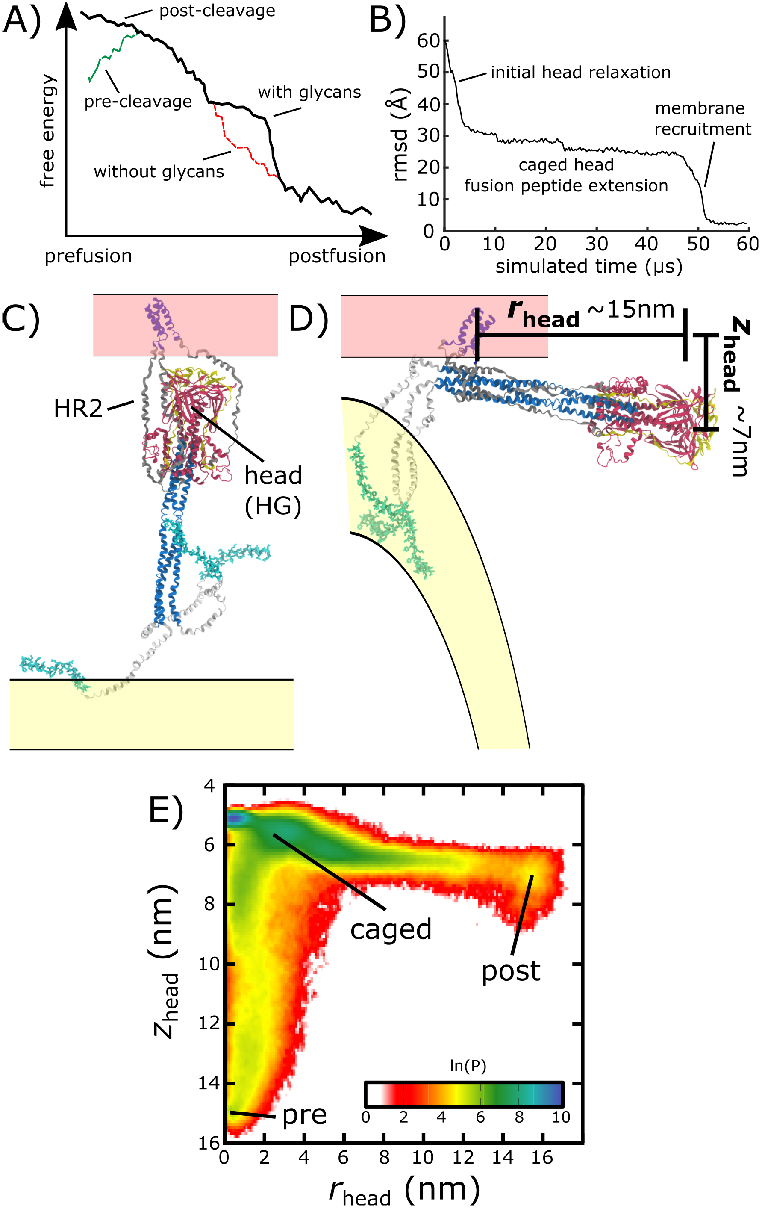
Simulating Spike-protein-mediated membrane fusion. Simulations with an all-atom structure-based model (*20, 21*) allow for transitions between prefusion and postfusion configurations to be observed. A) Schematic representation of the energetics in the structure-based model. The postfusion configuration was defined as the global potential energy minimum. The pre-cleavage state (green) is assumed to be stable, where cleavage and release of S1 leads to an unstable prefusion configuration (black). While, in the employed model, stabilizing energetic terms favor the postfusion configuration, steric interactions between the protein and glycans may impede the motion (black vs. red). B) Representative simulation (1 of 1000) of the pre-to-post transition. Spatial rmsd from the post configuration (excluding TM, CR and FPs), as a function of time. The simulation included explicit glycans, as well as an effective viral membrane potential. C) After an initial relaxation phase (panel B), the head (red) appears to become caged by the HR2 strands (gray), allowing it to sample configurations near the viral membrane (pink). While the host membrane (yellow) was not included in the simulations, it is depicted for illustrative purposes. D) After reaching the caged ensemble, the head escapes and the HR1-HR2 superhelix assembles. The position of the head group, relative to the TM region (blue), is described by the cylindrical coordinates *r*_head_ and *z*_head_. The origin is defined as the geometric center of TM, and the cylindrical axis is perpendicular to the viral membrane. See methods for details. E) Probability distribution of simulated events (with glycans) reveals an obligatory cage-like intermediate.

To investigate the dynamics of Spike-protein-mediated membrane fusion, we simulated thousands of transitions between the prefusion and postfusion conformations of S2 (Movie S1). To describe the rearrangements, we considered the distance between HG and the viral membrane (*z*_head_), as well as the displacement of HG parallel to the membrane (*r*_head_; Fig. 2D). The probability distribution as a function of *r*_head_ and *z*_head_ (Fig. 2E) shows a clear ordering of HG rearrangements. Each simulation was started from the prefusion conformation (*r*_head_ = 0, *z*_head_ *≈* 15nm). From there, the head moves towards the viral membrane (i.e. decreasing values of *z*_head_), and the HR2 strands appear to enclose HG. In this “caged” ensemble, the long axis of HG remains roughly perpendicular to the membrane (Fig. S2D). During initial relaxation of HG, the fusion peptides simultaneously extend toward the host membrane (Fig. 2C). After relaxation of HG, it then rotates away from its vertical orientation (increasing values of *r*_head_; Fig. 2D) by passing between two of the HR2 strands. As the head rotates (Fig. S2E-G), the FP and CR regions are drawn toward the viral membrane. The simulations were terminated when all non-CR and non-FP residues adopted their postfusion orientations. Since the CR and FP regions were not resolved in the postfusion structure (*30*), the simulations describe formation of all experimentally-resolved structural elements.

In simulations, the ordering of conformational events is robust to the presence of glycans. When glycans were explicitly included, there were only minor differences in the range of HG configurations that are sampled (Figs. 2D vs. S3). In both cases, HG initially relaxes towards the viral membrane before rotating towards the host.

### Glycans induce a long-lived sterically-caged intermediate

We find that glycans can reduce the kinetics of HG rearrangements by introducing a dynamic steric cage that confines HG to a position near the viral membrane (Fig. 3A). This caging process gives rise to prolonged sampling of an intermediate (Fig. 2B; 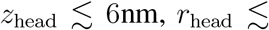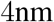) in which the long axis of HG is roughly perpendicular to the viral membrane (Fig. S2). The lifetime of the caged intermediate is given by *τ*_cage_ = *τ*_exit_ *− τ*_enter_ (Fig. 3B). *τ*_enter_ is defined as the time at which the assembly enters the intermediate (i.e. when *z*_head_ first decreases below 6.5 nm). *τ*_exit_ is the time at which *r*_head_ first exceeds 5 nm, indicating the head has been displaced outside of the cage-like formation (Fig. 2D). For the representative trajectory show in Fig. 3B, *τ*_cage_ is roughly 37*μs*. See methods for estimation of time units in this model.

**Figure 3:**
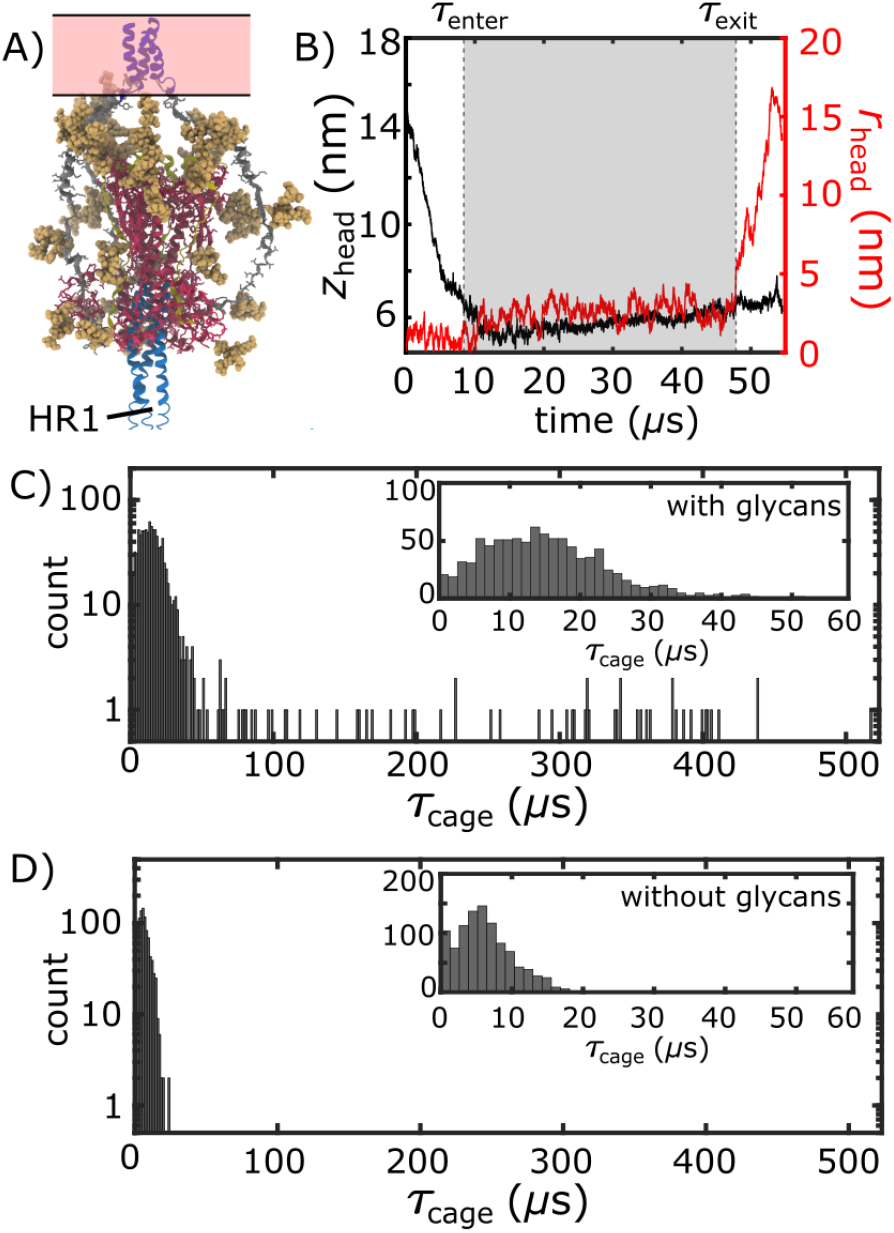
Glycan-induced caging of the head domain. Glycans impede head rearrangement by introducing a steric cage. A) Snapshot from the caged ensemble illustrates the high density of glycans surrounding the head. B) To define the duration of each caging event (*τ*_cage_ = *τ*_exit_ *τ*_enter_), we measured *z*_head_ and *r*_head_ (Fig. 2D). Based on the 2D probability distribution (Fig. 2E), the system was defined as entering the cage when *z*_head_ first drops below 6.5 nm: *τ*_enter_. *τ*_exit_ is the time at which the head moves laterally, relative to the trans-membrane region (*r*_head_ *>* 5 nm). C) Distribution of *τ*_cage_ values when glycans are present. There is an extended tail at large values (100 − 500*μs*). D) When glycans are absent, the *τ*_cage_ values are narrowly distributed around short timescales.

The lifetime of the caged intermediate strongly depends on the presence of glycans. For the glycan-free system, *τ*_cage_ values were narrowly distributed, where 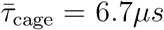 (Fig. 3D). When glycans are present, the distribution has a tail that extends to much larger values (100-500 *μs*; Fig. 3C), and 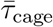 increases nearly 5-fold (29.7 *μs*). To isolate the origins of this effect, we repeated our simulations with subsets of glycans present. In one set of simulations, only glycans on HG were included, while the other set only included glycans on HR2 (Fig. S4). Interestingly, the HG-glycan model exhibited timescales that were comparable to those obtained for the fully glycosylated system. In addition, the HR2-glycan model yielded timescales that were comparable to those obtained when S2 is not glycosylated (Fig. S4). These comparisons reveal that the glycan-associated increase in excluded volume of HG is a dominant factor that determines the kinetics of interconversion between prefusion and postfusion conformations.

It is important to emphasize that the apparent glycan-dependent kinetics may be fully attributed to steric effects. That is, while the protein energetics were explicitly defined to favor the postfusion conformation, glycans were not assigned energetically-preferred conformations. Rather, the potential energy of the glycans only ensured that stereochemistry and excluded volume were preserved. In addition, the excluded volume interaction was purely repulsive. Thus, the observed reduction in rate for HG motion is due solely to the excluded volume of the glycans, and not the formation of stabilizing interactions. Finally, since glycans are likely to exhibit a degree of attraction under cellular conditions, the predicted glycan-induced reduction in kinetics should represent a lower-bound on the influence of glycosylation.

### Glycan cage promotes host membrane capture

The simulated trajectories suggest that glycan-associated attenuation of head rearrangements can facilitate host membrane recruitment and fusion. As described above, we find that the steric composition of the glycans introduce a highly crowded environment, which can transiently cage the HG domain (Fig. 3A). We additionally find that initial relaxation of HG is rapid (Fig. 2A), where caging introduces a pause that allows the HR1, FP and CR regions to sample extended configurations. To describe structure formation of the HR1 region, we calculated the fraction of postfusion-specific contacts that are formed as a function of time, *Q*_HR1_. Calculating the number of “native” contacts formed is motivated by protein folding studies, which have shown it to be a reliable measure of structure formation (*31*). When glycans are absent, HG frequently exits the cage prior to reaching the postfusion structure of HR1 (*Q*_HR1_ = 1200 *−* 1300; Fig. 4B). In contrast, when glycans are present, HR1 is typically fully-formed (*Q*_HR1_ *>* 1400) before HG exits the steric cage (Fig. 4A). By caging HG in a position that is perpendicular to the viral membrane (Fig. 3A), glycans help ensure that the newly-assembled HR1 helical coil remains directed towards the host membrane. This orientation of HR1 may serve to facilitate host membrane capture by the FP and CR regions.

**Figure 4:**
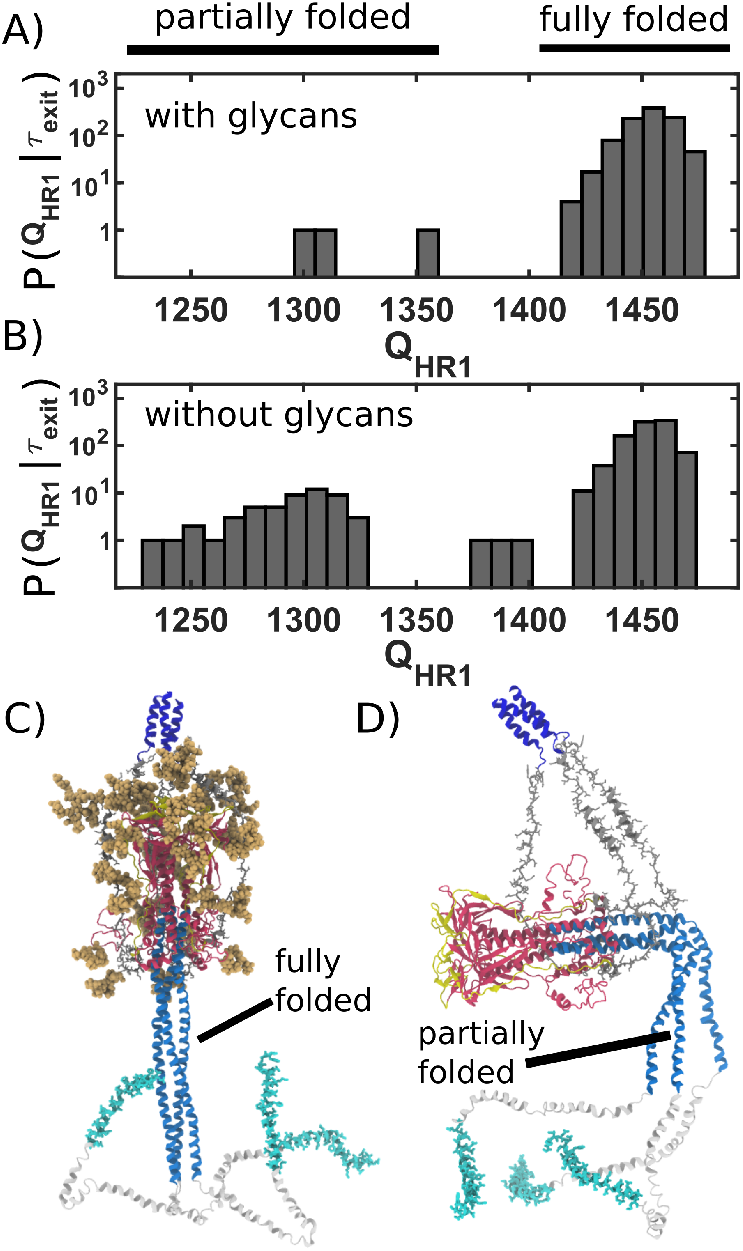
Caging of head allows for extension of HR1 helix. A) Distribution of *Q*_HR1_ (number of postfusion-specific HR1 contacts) values when glycans are present. Distribution describes the first frame in each simulations for which the head is outside of the steric c age. In all but 3 simulations, nearly all HR1 contacts (*>*1420 of 1489) are formed upon exit of the cage. B) Distribution when glycans are not included. When glycans are absent, it is common that HR1 is not completely formed (i.e. *Q*_HR1_ *<*1350) prior to HG escape. C) Representative snapshot of a caged structure in which HR1 is fully formed and extended toward the host. D) Representative snapshot of a glycan-free case where the head escapes prior to fully forming HR1. As a result, HR1 can adopt bent configurations.

To quantify the likelihood that the Spike protein will associate with and recruit the host, we considered the extension of each fusion peptide from the viral membrane. For this, we first defined a putative host-membrane distance, *d*_host_, which was set to discrete values (22-38 nm). We then determined whether the distance between the viral membrane and FP (*d*_FP_, Fig. 5A) exceeded *d*_host_ for each of the three FP tails in the assembly (Fig. 5B). *P*_capture_ was then defined as the probability that at least one FP extends to the host membrane (Fig. 5C). We use the notation *P*_capture_, since one expects that the extension of the FPs will be correlated with the probability that the Spike protein successfully captures the host cell.

**Figure 5:**
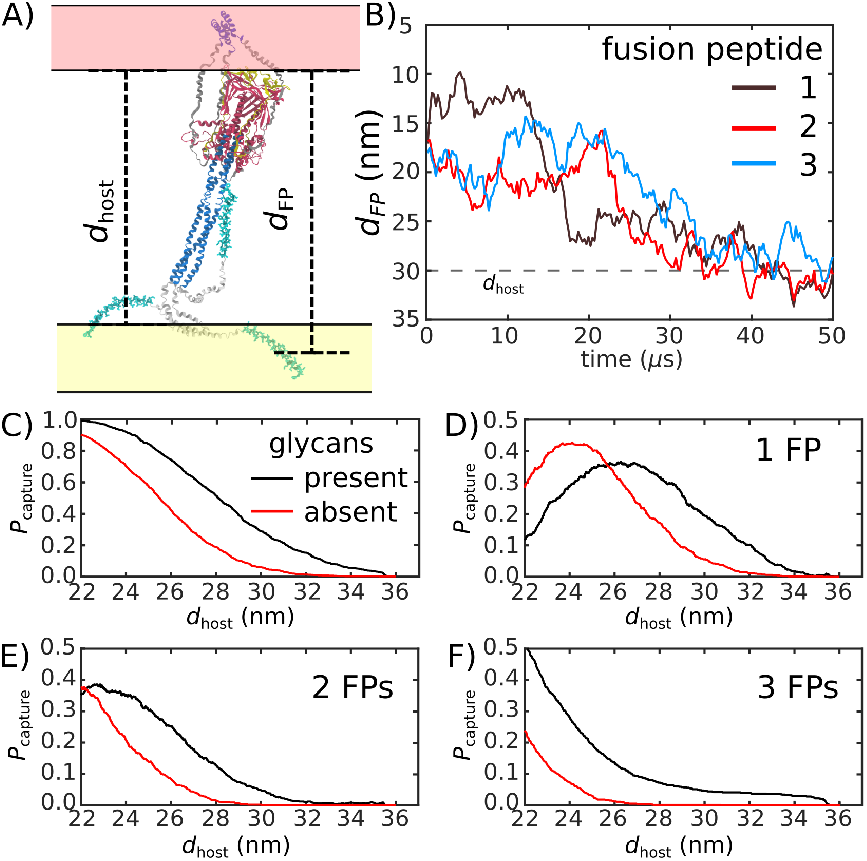
Glycans promote host capture. A) Snapshot of the Spike protein with the head domain in a caged configuration. Caging allows the fusion peptide tails to extend toward and engage the host membrane. *d*_FP_ is the distance of the center of mass of each fusion peptide from the viral membrane surface. To calculate the probability of host capture, different values of the virion-host distance (*d*_host_) were considered. B) Representative simulated trajectory, showing *d*_FP_ for each of the fusion peptides in a single S2 subunit. For reference, a *d*_host_ value of 30 nm is indicated by a dashed line. C) The probability of membrane fusion is expected to be proportional to the probability that at least one tail extends to the host membrane (*d*_FP_ *> d*_host_). There is a higher probability of extending to larger *d*_host_ values when glycans are present (black vs. red curves). This is due to the glycan-induced delay of head rotation (Fig. 3), which ensures the HR1 helix remains directed towards the host as the FPs sample extended configurations. D-F) The probability that 1, 2 or 3 FPs exceed *d*_host_. In all cases, the presence of glycans shifts the distribution to larger values of *d*_host_, indicating an increased probability of capturing the host. This reveals a critical role for glycans during cell invasion.

We find that glycosylation of S2 significantly increases the probability that a FP will extend to the host membrane (Fig. 5C). We further partitioned the capture distributions by calculating the probability that exactly 1, 2 or 3 FPs cross the host membrane (Fig. 5D-F). Interestingly, we find that all distributions exhibit dramatic differences for 26 *< d*_host_ *<* 34 nm. In the absence of glycans, the probability of associating 3 FPs is nearly 0 in this range, while the probability increases to *~* 0.1 when glycans are present (Fig. 5F). Similarly, the probability that exactly 2 FPs will cross the host membrane is *~* 0 for *d*_host_ *>* 28 nm, when glycans are absent. The probability then increases to *~* 0.1 when glycans are present (Fig. 5E). Finally, for 26 *< d*_host_ *<* 33, the probability that one FP will cross the host membrane is increased by *~*0.2 when glycans are present (Fig. 5D).

The glycan-dependent probability of host-membrane association suggests several features of the fusion process. Cryotomography imaging has revealed that the virus-host inter-membrane distance is approximately 30nm during infection (*13*). Based on this, our simulations indicate that, if the Spike protein were not glycosylated, it is most likely that none of the FPs would associate with the host. Since Spike protein rearrangements are irreversible, these failed attempts would represent lost opportunities to infect the cell. Further, when glycans are absent, there is a marginal probability that only one or two FPs would reach the membrane, where the other FPs would likely transition directly to their postfusion orientations without engaging the host. Such a process has been described as a “cooperative” mechanism in the Hemagglutinin A fusion protein (*32*). However, when glycans are present, there is a non-negligible probability that three FPs will reach the host membrane. In that case, the dynamics may also involve the so-called “sequential” mechanism of fusion (*32*). Together, these observations demonstrate how the steric contributions of glycans is critical to the mechanism and likelihood of cell invasion by SARS-CoV-2.

## Discussion

The ongoing COVID-19 pandemic requires the rapid identification of molecular factors that enable infection. A necessary step during infection involves virus/cell membrane fusion, which is mediated by a major conformational change of the Spike protein. Here, we propose a mechanism where, after cleavage and dissociation of S1, sufficient time has to be made available for the fusion peptides to reach the cell membrane, before the conformational change in S2 can complete. Using all-atom models with simplified energetics, we have shown how the steric composition of post-translational modifications may introduce the delay necessary for such a mechanism to be utilized. This glycan-induced pause appears to allow for an extended window during which the fusion peptides may search for the host cell (Fig. 6). In simulations that did not include glycans, the Spike protein was most likely to adopt the postfusion configuration without extending the fusion peptides towards the host. Thus, in the glycan-free case, the protein can bypass the sterically-caged intermediate, leading to failed attempts to capture the host cell. These findings suggest that the precise glycan composition is a critical factor that determines transmissibility of SARS-CoV-2.

**Figure 6:**
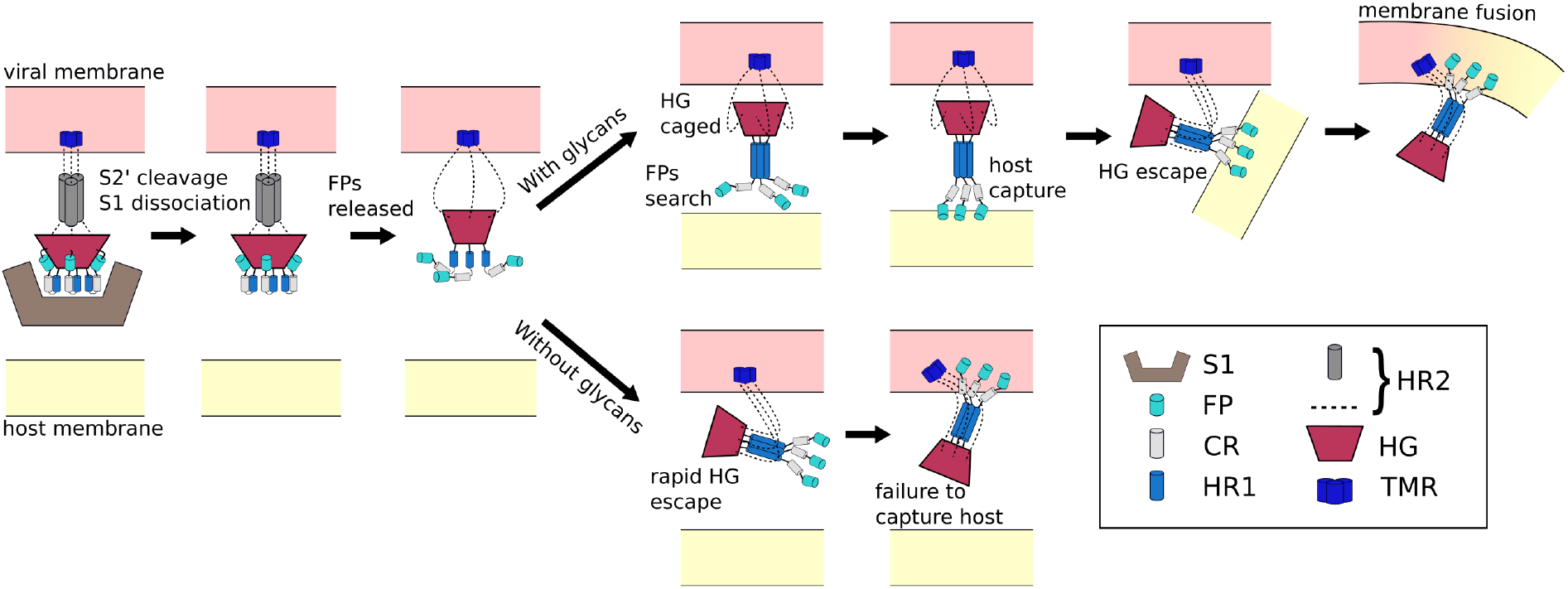
Schematic of transition mechanism of the Spike protein. Initial activation of the Spike protein (left) is associated with release of S1, which is triggered by cleavage at the S2’ site and ACE2 receptor binding. When glycans are present (top), HG will enter a caged ensemble where the FPs search for, and capture, the host membrane. HG can then escape the cage, which draws the viral and host membranes together and leads to fusion. In the absence of glycans (bottom), HG can bypass the caged ensemble, resulting in a failed attempt to recruit the host.

In addition to providing immediate insights into the influence of glycans on Spike protein dynamics, the current simulations establish a foundation for investigating other factors that may influence cell invasion. For example, the presented models may be extended to account for electrostatic and solvation effects. In addition, virtual mutations (*29*) could be used to quantify the relative importance of different structural elements. Such approaches could identify how to more effectively disrupt the various steps involved with membrane fusion, which could ultimately serve to neutralize the virus. With ongoing advances in high-performance computing, combined with the relatively low computational cost associated with these models, many variations may be explored in the coming months that will help elucidate the full range of factors that control this deadly pathogen.

## Materials and Methods

### Spike Protein Modeling

Since complete structures of the full-length SARS-CoV-2 Spike protein have not been resolved experimentally, for either the prefusion or postfusion states, structural modeling steps were applied prior to performing simulations. For this, we used a cryo-EM structure of the prefusion assembly (PDB ID: 6VXX (*2*)), which lacks residues 828-853 and 1148-1233 (found in HG, and the HR2 and TM regions). For the postfusion state we used a structural model of the SARS-CoV-1 system (PDB ID: 6M3W (*30*)) as a template for constructing a homology model of the SARS-CoV-2 system. However, the postfusion model was lacking residues 772-918 and 1197-1233 (FP, CR and TM). In addition, since available structures only partially resolved the base of each glycans, we constructed structural models of the complete glycans for both states (pre and post). For the prefusion structure, models of the TM and HR2 regions were constructed using the homology modeling webtool of SWISS-MODEL (*33*). Consistent with the study of Casalino et al (*7*), both regions were modeled as coiled coils, where the sequence was assigned Uniprot (*34*) sequence P0DTC2-1. This was accomplished with the automodel module with special restrains to preserve symmetry of Modeller 9.24 (*35*). The resulting model was three-fold symmetric, where the RMSD between monomers was less than 1 Å. For the postfusion structure, missing residues in FP, CR and TM were modeled as helical regions, using the automodel module of Modeller 9.24. CR and FP were modeled as coiled coils that are connected by short disorder loops. To define the CR and FP regions, homology models were constructed based the structure of a coiled coil template (PDB ID: 2WPQ (*36*)). Glycans were added to both structural models using the Glycan Reader Charmm server (*37*). The same glycan composition was used as in other recent studies (*6, 7*). A complete list of modeled glycans can be found in Table S1.

### All-atom structure-based model

All simulations employed an all-atom structure-based model to describe the Spike protein, with additional restraints imposed on the TM region, as well as an effective viral membrane potential. To describe the energetics of the protein, a structure based model was constructed based on the postfusion model, using the default parameters in SMOG 2 (described in Ref. (*21*)). Several modifications were introduced to the force field, as described below. Non-default parameters were assigned for bond lengths and angles, as well as planar dihedrals. Rather than using the values found in the cryo-EM structure, bond lengths and angles were given the values found in the Amber03 force field (*38*). For planar dihedrals, a cosine function of periodicity 2 with minima at 0 and 180° was applied. The strengths of all interactions were consistent with earlier implementations of the structure-based model (*21*). That is, all non-planar dihedral angles were assigned 1-3 dihedral potentials with minima corresponding to the postfusion conformation. Stabilizing 6-12 interactions were introduced for all atom pairs that are in contact in the postfusion conformation, with minima at distances corresponding to the postfusion conformation. Contacts were identified using the Shadow algorithm (*39*). A complete description of this variant of the model is described in Ref. (*40*). The model is available for download at https://smog-server.org (SMOG2 force field repository name: SBM AA-amber-bonds). To ensure that the TM region remains in a helical bundle arrangement, contacts in the TM region were replaced by harmonic interactions, with distances taken from the prefusion conformation. To mimic the presence of a viral membrane, a flat-bottom potential show in Eq. 1 was imposed on the TM region to limit movement inside the putative membrane region. Also, to avoid non-TM residues from crossing the effective membrane, an inverted harmonic flat bottom potential was applied to center of mass of HG. The potential was set to 0 at *z*_head_ = 2 and the harmonic constant was set to 2 reduced energy units per nm).

### MD simulations

All simulations were performed using the GROMACS software package (v2020.2) (*41,42*) with source code modifications to implement the Gaussian-based flat bottom potential (Eq. 1). Input files for Gromacs were generated using the SMOG 2 software package (*21*), while additional in-house scripts were used to subsequently modify the force field. Simulations of four different molecular systems were performed: glycan-free, fully-glycosylated, HR2-glycosylated and HG-glycosylated. 1000 transitions between prefusion and postfusion structures were simulated for each system (4000 simulations, in total). Each system was first energy minimized using steepest descent energy minimization. Simulations were then performed using Langevin Dynamics protocols, with a reduced temperature of 0.58 (70 Gromacs units). In preliminary simulations, it was found that the assembly begins to unfold at a temperature of around 0.8. A timestep of 0.002 was used, and each simulation was continued until *r*_head_ reached a value greater than 8 nm, which indicated that HG had escaped from the HR1 cage. To estimate the effective simulated timescale, we use the conversion factor of 1 reduced unit being equivalent to 1 ns (*43*), which was previously obtained based on the comparison of diffusion coefficients in a SMOG model and explicit-solvent simulations.

### Structural metrics

The following coordinates were used to describe the global rearrangement of the Spike protein:

- *z*_head_: To calculate *z*_head_, the vector between the centers of mass of TM (residues 1203-1233) and HG (residues 1033-1129) was calculated and then decomposed into components that are perpendicular and parallel to the membrane plane. *z*_head_ is the component that is perpendicular to the plane.
- *r*_head_: To calculate *r*_head_, the vector between the centers of mass of TM (residues 1203-1233) and HG (residues 1033-1129) was calculated and then decomposed into components that are perpendicular and parallel to the membrane plane. *r*_head_ is the component that is parallel to the plane.
- *θ*: Angle formed between the first principal axis of HG (residues 1033-1129) and the vector normal to the membrane. A value of 0 indicates that the HG is perpendicular to the viral membrane.
- *Q*_HR1_: Number of postfusion-specific contacts formed (within 1.2 times the distance in the post-fusion conformation).

#### Membrane flat-bottom potential

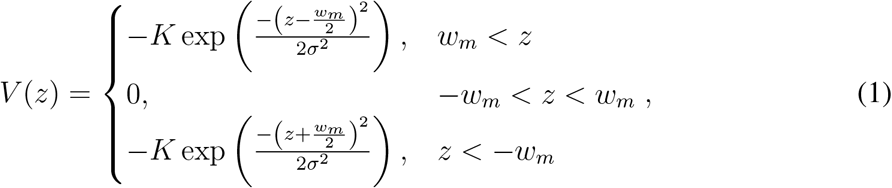

where *z* is the distance of the center of mass of TM from the center of the effective membrane. *w*_*m*_ represents the membrane width (5nm), and the depth of the potential *K* was set to 2 reduced energy units.

## Acknowledgments

JNO was supported by the NSF grants CHE-1614101 and PHY-1522550 and by the Welch Foundation (Grant C-1792). PCW was supported by NSF grant MCB-1915843. Work the Center for Theoretical Biological Physics was also supported by the NSF (Grant PHY-2019745). JNO is a CPRIT Scholar in Cancer Research. We would also like to acknowledge generous support from the Northeastern University Discovery cluster and NU Research Computing staff.

## Supplementary materials

Figs. S1 to S4

Tables S1

Movie S1

