## Supplementary Information for "Sterically-Confined Rearrangements of SARS-CoV-2 Spike Protein Control Cell Invasion"

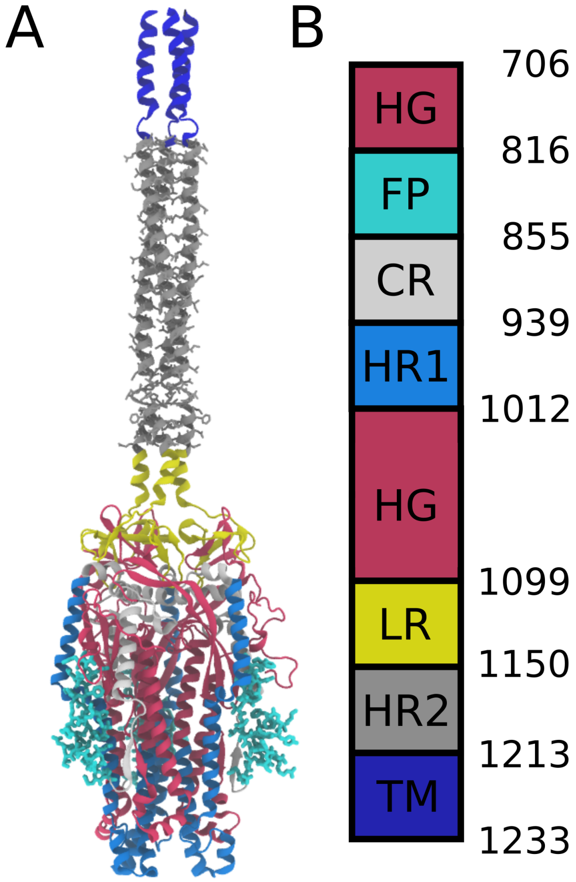


Fig. S1: **Definitions of domains within the S2 protein.** A) Prefusion S2 subunit structure of the Spike protein. B) Sequence range of the Head Group (HG), Fusion Peptide (FP), Connecting Region (CR), Heptad Repeat 1 (HR1), Linker Region (LR), Heptad Repeat 2 (HR2), and Transmembrane Region (TM).

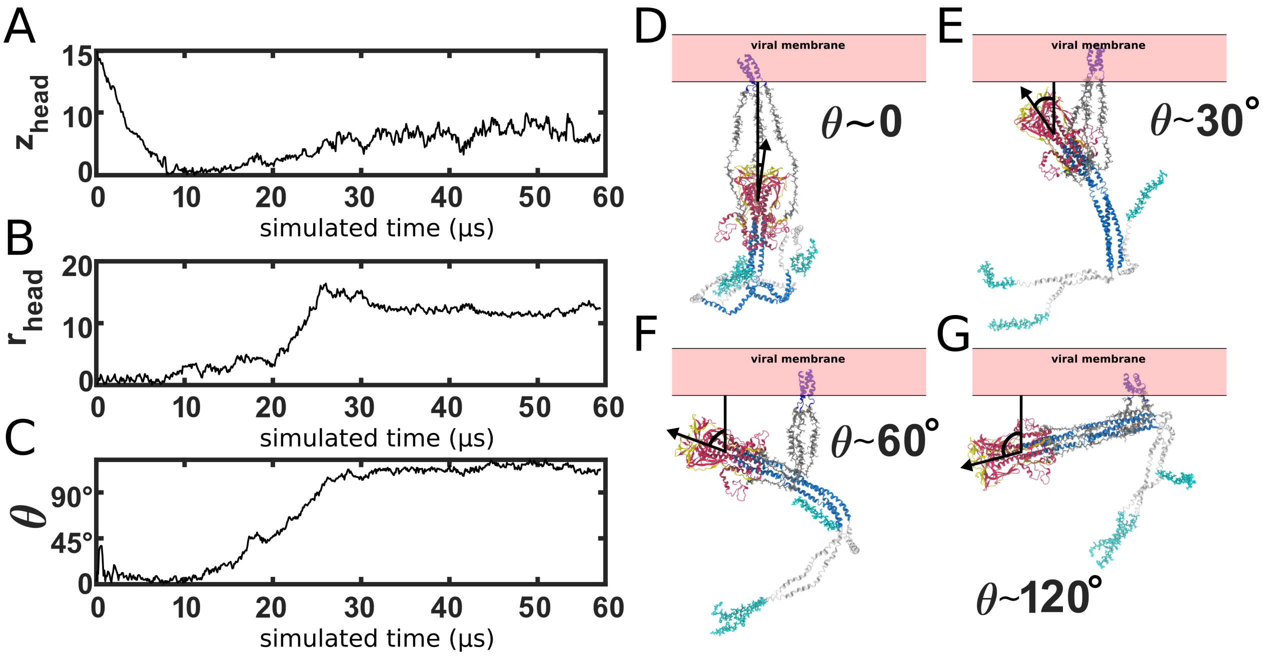


Fig. S2: **HG rotation** A-C) Single time trace of $z_{\mathrm{head}}$, $r_{\mathrm{head}}$ and the HG principal axis polar angle, $\theta$. D-G) Snapshots of the orientation of HG, relative to the membrane. During the prefusion-to-postfusion transition, the head rotates from an orientation in which it is pointing towards the membrane, to an orientation where it is pointing away. Structural snapshots illustrate various orientations during the transition.

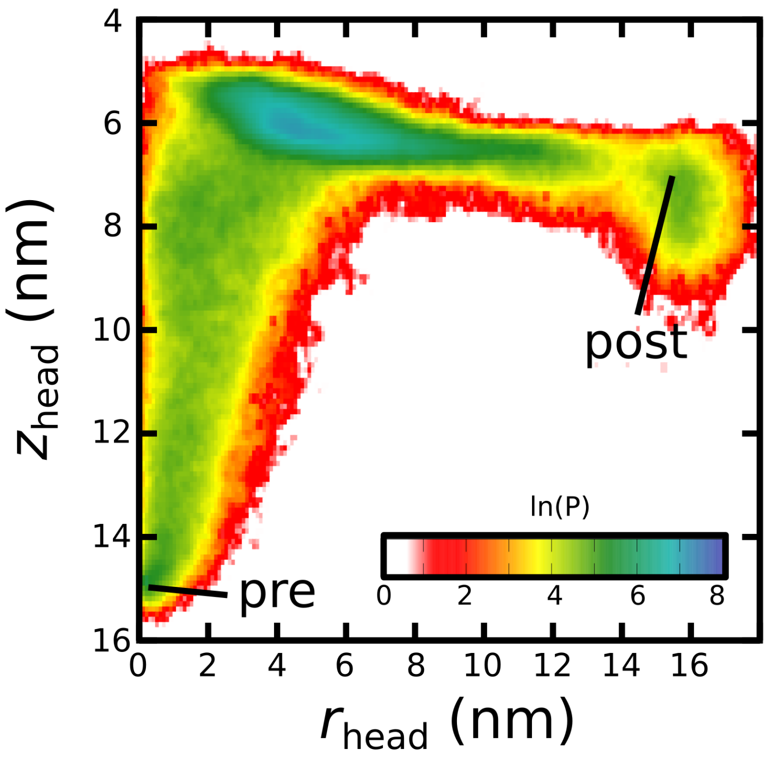


Fig. S3: **Probability distribution when glycans are absent** Distribution calculated from 1000 independent simulations without glycans.

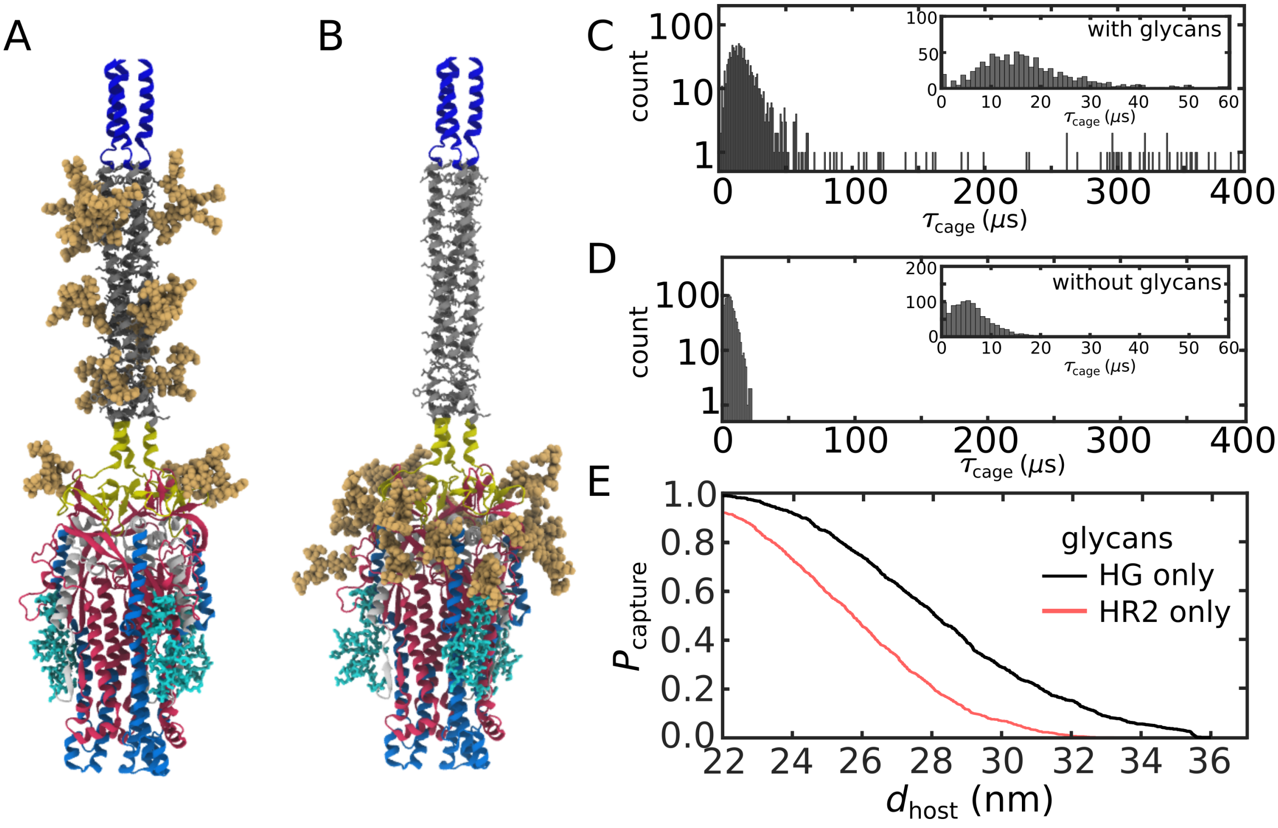


Fig. S4: **Relative influence of HR2 and HG glycans.** A) Structural model with only glycans shown on HG. B) Structural model with only HR2 glycans present. C) Distributions of timescales with HG-only glycans. D) Distribution with HR2-glycans. E) Probability that $d_{FP} > d_{host}$ for at least one FP. There is a higher probability of extending to $d_{host}$ when HG-only glycans are present than when HR2-only glycans are present (black vs. red curves)

Tab. S1: **N-glycan listing** Complete list of N-glycans included in the simulations.

| | |
|---|---|
| N706 | aDMan(1→6)[aDMan(1→3)]aDMan(1→6)[aDMan(1→3)] bDMan(1→4)bDGlcNAc(1→4)bDGlcNAc(1→) |
| N717 | aDMan(1→6)[aDMan(1→3)]aDMan(1→6)[aDMan(1→2)aDMan(1→3)] bDMan(1→4)bDGlcNAc(1→4)bDGlcNAc(1→) |
| N801 | aDMan(1→6)[aDMan(1→3)]aDMan(1→6)[aDMan(1→3)] bDMan(1→4)bDGlcNAc(1→4)bDGlcNAc(1→) |
| N1074 | aDMan(1→6)[aDMan(1→3)]aDMan(1→6)[aDMan(1→3)] bDMan(1→4)bDGlcNAc(1→4)bDGlcNAc(1→) |
| N1098 | aDNeu5Ac(2→6)bDGal(1→4)bDGlcNAc(1→2)aDMan(1→3)[aDMan(1→6) [aDMan(1→3)]aDMan(1→6)]bDMan(1→4)bDGlcNAc(1→4)bDGlcNAc(1→) |
| N1134 | bDGlcNAc(1→2)aDMan(1→6)[bDGlcNAc(1→2)aDMan(1→3)]bDMan(1→4) bDGlcNAc(1→4)[aLFuc(1→6)]bDGlcNAc(1→) |
| N1158 | bDGlcNAc(1→2)aDMan(1→6)[bDGlcNAc(1→2)aDMan(1→3)]bDMan(1→4) bDGlcNAc(1→4)bDGlcNAc(1→) |
| N1173 | bDGlcNAc(1→6)[bDGlcNAc(1→2)]aDMan(1→6)[bDGlcNAc(1→4)[bDGlcNAc(1→2)] aDMan(1→3)]bDMan(1→4)bDGlcNAc(1→4)[aLFuc(1→6)]bDGlcNAc(1→) |
| N1198 | aDNeu5Ac(2→6)bDGal(1→4)bDGlcNAc(1→6)[bDGal(1→4)bDGlcNAc(1→2)] aDMan(1→6)[bDGal(1→4)bDGlcNAc(1→4)[bDGal(1→4)bDGlcNAc(1→2)] aDMan(1→3)]bDMan(1→4)bDGlcNAc(1→4)[aLFuc(1→6)]bDGlcNAc(1→) |
